# A general goodness-of-fit test for survival analysis

**DOI:** 10.1101/104406

**Authors:** Michael Holton Price, James Holland Jones

## Abstract

Existing goodness-of-fit tests for survival data are either exclusively graphical in nature or only test specific model assumptions, such as the proportional hazards assumption. We describe a flexible, parameter-free goodness-of-fit test that provides a simple numerical assessment of a model’s suitability regardless of the structure of the underlying model. Intuitively, the goodness-of-fit test utilizes the fact that for a good model early event occurrence is predicted to be just as likely as late event occurrence, whereas a bad model has a bias towards early or late events. Formally, the goodness-of-fit test is based on a novel generalized Martingale residual which we call the martingale survival residual. The martingale survival residual has a uniform probability density function defined on the interval −0.5 to +0.5 if censoring is either absent or accounted for as one outcome in a competing hazards framework. For a good model, the set of calculated residuals is statistically indistinguishable from the uniform distribution, which is tested using the Kolmogorov-Smirnov statistic.

## 1 Introduction

Assessing the goodness-of-fit of survival models presents challenges not encountered with conventional regression models. In a conventional statistical analysis, the dependent variable or variables being modeled are typically also directly observed. For example, the heights of a sample of individuals are both directly measured and predicted with a model that can account for covariates such as age and gender. In contrast, in survival analysis the timing of event is observed (e.g., age at death) but hazard is predicted. This complicates the definition and use of common tools for assessing goodness-of-fit, such as data residuals. An additional complication is censoring, which is common in survival data and, e.g., skews the distribution of martingale residuals. Due to these complications, no goodness-of-fit test exists that is simultaneously general, robust, and straightforward to use. In this article, we describe a simple, flexible, and parameter-free goodness-of-fit test based on a novel data residual, the martingale survival residual.

The martingale survival residual belongs to a class of generalized martingale residuals that includes the conventional martingale residual (Barlow & Prentice 1988). The conventional martingale residual equals the difference between the number of occurrences of an event at some stopping time and the estimated cumulative hazard of that event up to the stopping time (Equation 11). The value of the residual depends on whether the observation is censored. For events that occur only once for each subject (e.g., death in demographic models), it equals 1 minus the estimated cumulative hazard for non-censored observations and 0 minus the estimated cumulative hazard for censored observations. Because the cumulative hazard takes on values between 0 and ∞, the martingale residual takes on values between −∞ and 1.

We define a new generalized martingale residual, the martingale survival residual, which is based on the estimated survival and takes on values on a finite range, −0.5 to +0.5 (Equation 12). The theoretical probability density function of the martingale survival residual is the uniform distribution defined on the interval −0.5 to +0.5, so long as censoring is absent or is accounted for as one event in a competing hazards framework. Negative values of the martingale survival residual correspond to “early” events and positive values to “late” events. The uniform nature of the martingale survival residual’s probability density function makes it easier to interpret than the martingale residual and is the basis for the goodness-of-fit test described in this article.

Since the theoretical probability density function of the martingale survival residual is a uniform distribution, a good survival model will yield a set of martingale survival residuals that are statistically indistinguishable from the uniform distribution. In contrast, a bad model will be statistically distinguishable, which can be tested using well established techniques such as the Kolmogorov-Smirnov test or the Anderson-Darling test. In this article, we utilize the Kolmogorov-Smirnov test to implement a goodness-of-fit test, but suggest that future work should consider other tests.

The remainder of this article is structured thus: (1) Sections 2 and 3 present the theoretical material for the goodness-of-fit test; (2) Section 4 illustrates the effectiveness of our goodness-of-fit test using simulated data; and (3) Sections 5 and 6 provide a discussion and concluding remarks, including avenues for future work, such as how to accommodate data sets with terminal censoring – i.e., studies with a terminal end date when censoring is guaranteed to occur, which creates a discontinuity in the censoring hazard.

## 2 Residuals for event history analysis

### 2.1 Preliminaries

Let *N_i_*(*t*), *Y_i_*(*t*), and *Z_i_*(*t*) represent, respectively, the counting, risk, and covariate processes for observation *i*, so that 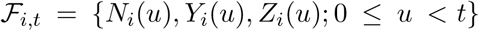 completely specifies the realized history (σ-algebra) of observation *i* up to time *t* (Barlow & Prentice 1988). Similarly, let *N_i_^C^* (*t*), *Y_i_^C^* (*t*), and *Z_i_^C^* (*t*) represent the counting, risk, and covariate processes for the generally hypothetical situation in which all outcomes are observed without censoring, so that {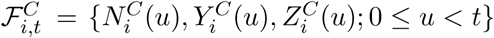 completely specifies the hypothetically realized history (σ-algebra) of observation *i* up to time *t*. This notation and terminology follow Aalen et al. (2008). *Y_i_* (*t*) and *Y_i_^C^* (*t*) are linked through the equation

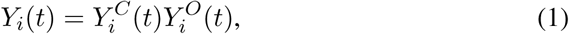
Where *Y_i_^O^*(*t*) is a left continuous binary censoring process which equals 1 if an individual is under observation immediately prior to time *t* and 0 otherwise. The intensity process (or hazard) of *N_i_^C^*(*t*) relative to the history 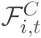 takes the form

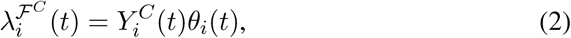

where the functions *θ_i_*(*t*) are assumed to be 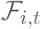-predictable. Assuming independent censoring, the intensity of *N_i_*(*t*) relative to the history 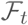 takes the form

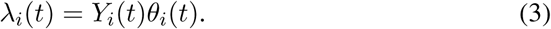

The assumption of independent censoring will be relaxed for the treatment of the comprehensive hazard function. Hence, it is not crucial to the goodness-of-fit test. However, if independent censoring is rejected, a suitable model for joint event and censoring hazards must be defined.

### 2.2 Generalized martingale residuals

Invoking the Doob-Meyer theorem, any counting process *N*(*t*) can be decomposed into a unique, predictable cumulative intensity process
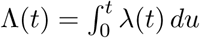, assumed to be absolutely continuous, and a zero mean martingale *M*(*t*),

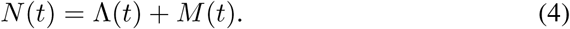

Because the transformation of a predictable process with respect to a zero mean martingale is also a zero mean martingale, Equation 4 can be recast to yield the generalized martingale residual corresponding to the predictable process *H*(*t*),

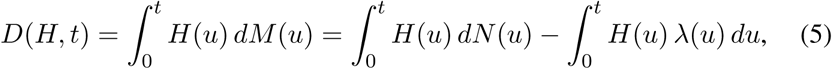

Where λ(*t*) = Λ′(*t*) and *D*(*H, t*) is a zero mean martingale by construction. The first integral on the right hand side of Equation 5 is a stochastic integral, which is equal to the sum of the values of *H* at all jump times of *N* up to *t*,

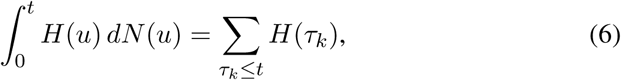
where *τ_k_* are the jump times of the counting process *N*. The second integral in Equation 5 is a conventional Riemann integral. In the general framework described by Barlow & Prentice (1988), for each predictable process *H_i_* associated with observation *i* there is both a corresponding actual data residual

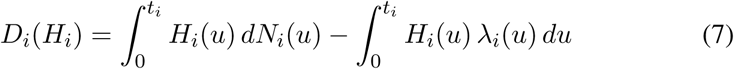
And a corresponding predicted data residual

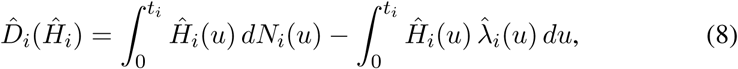
where *t_i_* is the stopping time of observation *i* and the hats distinguish predicted variables, functions, or functionals from actual ones. 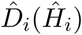 is what a researcher calculates and analyzes after modeling a data set of interest. In addition, actual and predicted data residuals can be defined for the generally hypothetical complete counting process *N_i_^C^*(*t*),

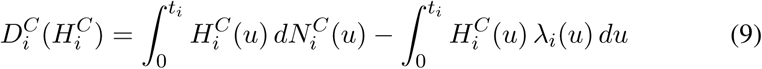
and

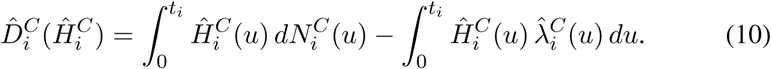

Censoring causes the expected probability density functions associated with 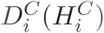
 and 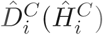
 to differ from those associated with *D_i_* (*H_i_*) and 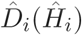.

### 2.3 The conventional martingale residual and martingale survival residual

The choice *Ĥ_i_* = 1 yields the conventional martingale residual, whereas the choice 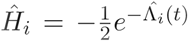 yields the martingale survival residual, which is a linear transformation of the predicted survival function, 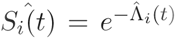
. The negative sign on the martingale survival residual makes it so that earlier exit times correspond to smaller residuals, which is more intuitive to interpret than the reverse. For the same reason, we will use the negative martingale residual with *Ĥ_i_* = −1 in this article rather than the conventional martingale residual.

We assume that the counting process *N*_*i*_(*t*) is capped at 1 since this is true for many applications of event history analysis and the assumption simplifies the explication. Generalizations that relax this assumption are straightforward. The formulas for the negative martingale residual and martingale survival residual are then, respectively,

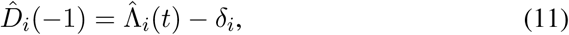
and

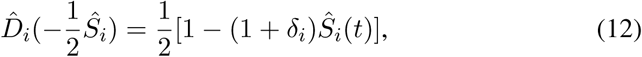
where *δ_i_* is 1 for observed events and 0 for censored outcomes. In the Appendix, we derive theoretical probability density functions for the actual data residuals corresponding to the predicted data residuals given by Equations 11 and 12 (i.e., dispensing with the hats). The probability density functions of the negative martingale residual and martingale survival residual with no censoring are

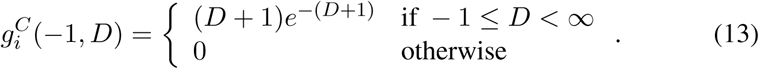
and

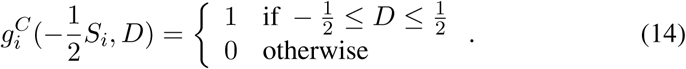

If censoring is present, the probability density functions depend on the functional form of the event and censoring intensities. For the special case of constant transition intensities for both event occurrence and censoring, the probability density functions of the negative martingale residual and martingale survival residual are

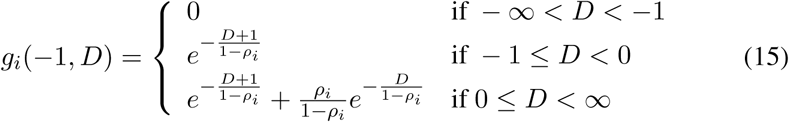
and

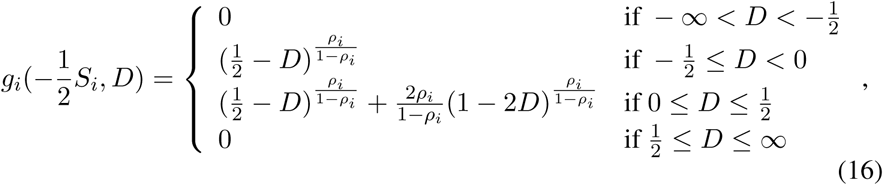

where *ρ_i_* = *c*_*i*_/(λ_*i*_ + *c*_*i*_) is the censoring ratio – that is, the probability that subject *i* is censored. Figures 1 and 2 plot the probability density functions for the negative martingale residual and martingale survival residual given the assumption of constant transition intensities (Equations 15 and 16, both with no censoring (which recovers the probability density functions in Equations 14 and 14) and with 25% censoring. The plots have a discontinuity at *D* = 0 due to censoring.

**Figure 1:**
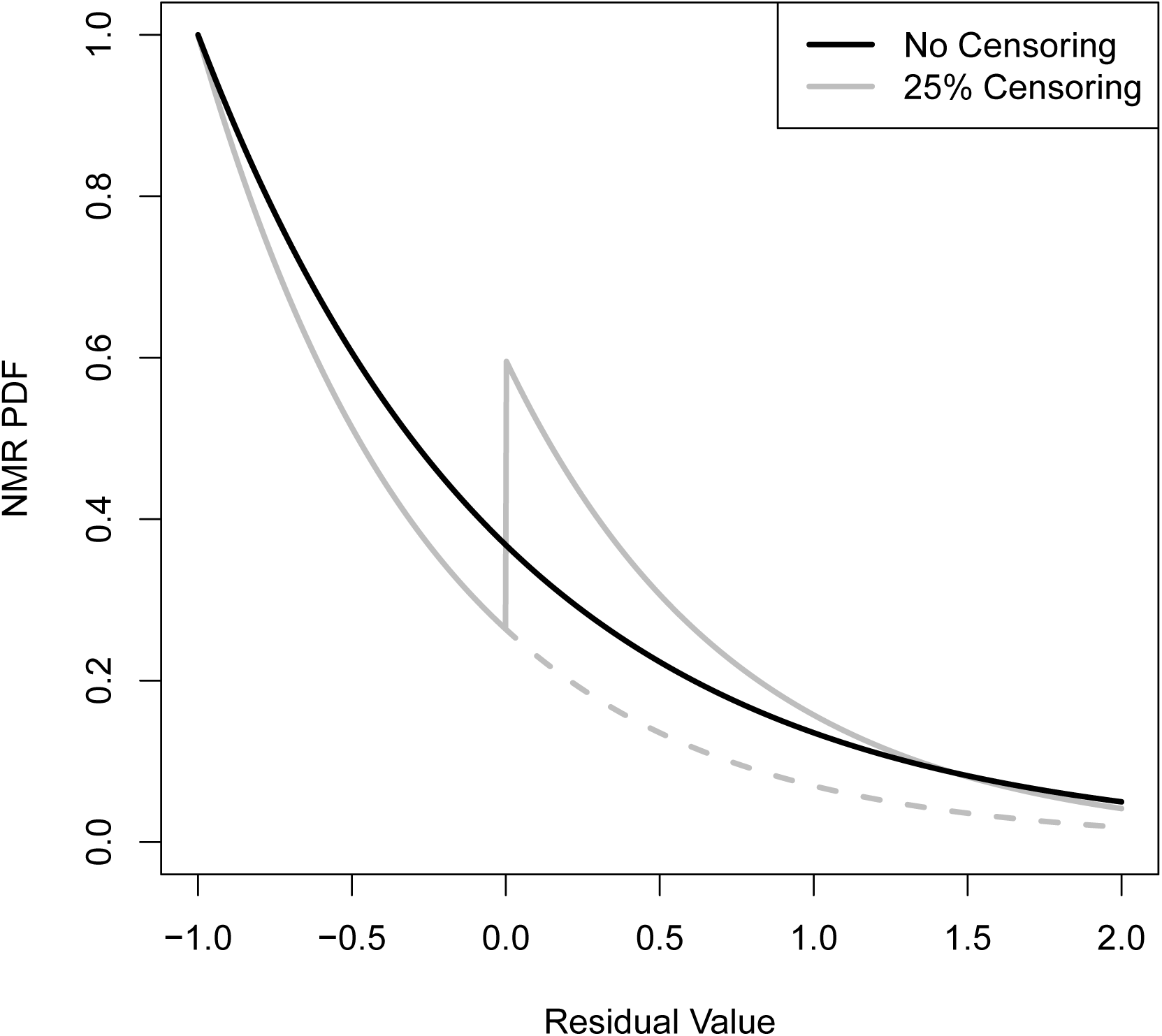
Expected probability density function for the negative martingale residual assuming constant event occurrence and censoring transition intensities for censoring percentages of 0% and 25%. Censoring only contributes for *D* ≥ 0. The dashed line shows the contribution of non-censored outcomes to the probability density function for the 25% censoring case. The difference between the solid and dashed lines is the contribution of censored outcomes.

**Figure 2:**
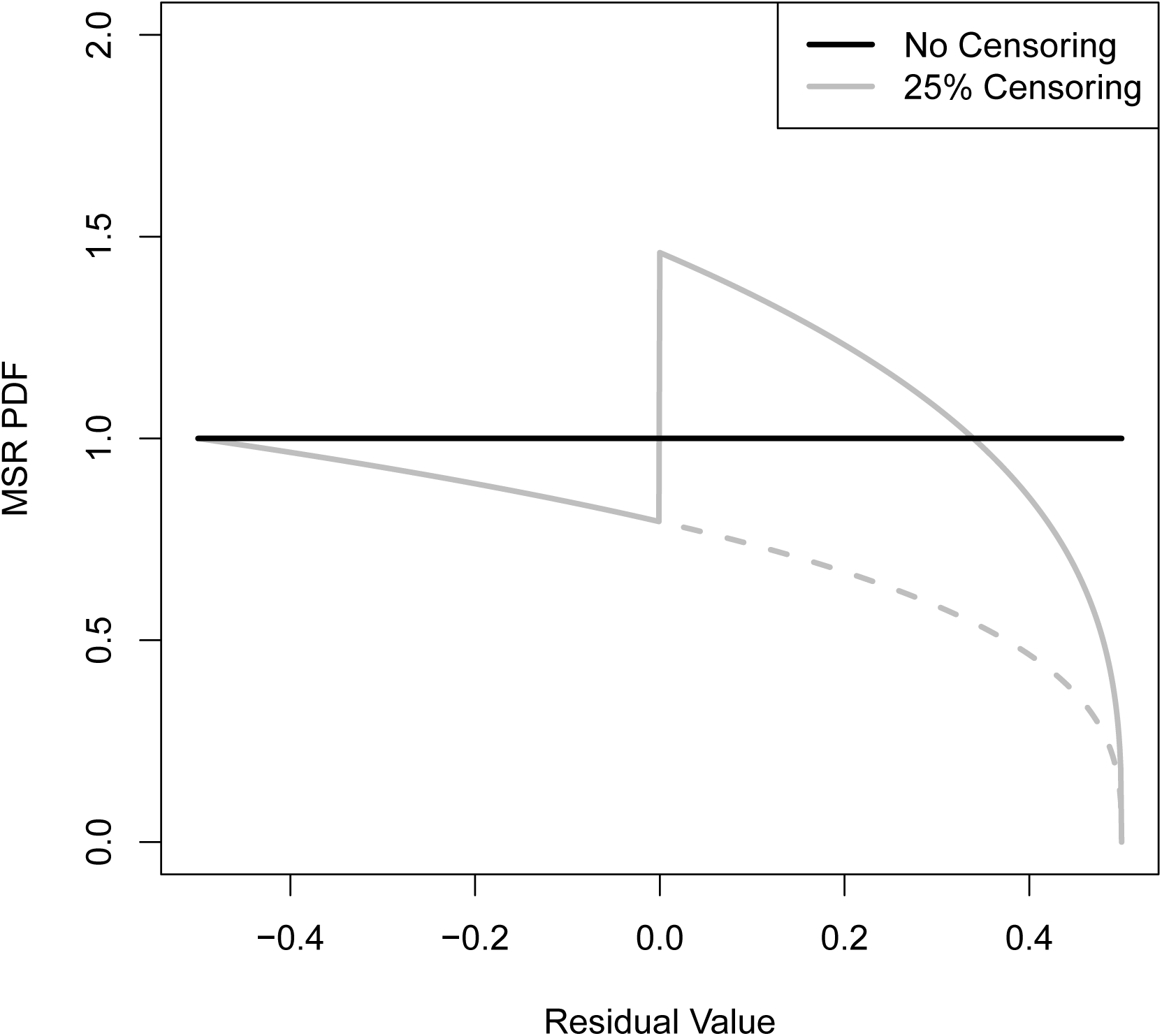
Expected probability density function for the martingale survival residual assuming constant event occurrence and censoring transition intensities for censoring percentages of 0% and 25%. Censoring only contributes for *D* ≥ 0. The dashed line shows the contribution of non-censored outcomes to the probability density function for the 25% censoring case. The difference between the solid and dashed lines is the contribution of censored outcomes.

In the absence of censoring – or if censoring is accounted for as one event in a competing events framework – the probability density function for the martingale survival residual is particularly simple: it is a uniform distribution defined on the interval −0.5 to +0.5. This has substantial intuitive appeal, we suggest, since a good model of event occurrence (i.e, a good model of the transition intensity 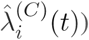 is equally likely to lead to early exits as late exits. Hence, the predicted survival at the transition time, *Ŝ_i_*, should be drawn from a uniform distribution. Extending this observation to a set of observations, a good model of transition intensities should yield a set of martingale survival residual residuals that is statistically indistinguishable from the uniform distribution. This is the basis for the goodness-of-fit test described in the next section.

## 3 A goodness-of-fit test based on the Kolmogorov-Smirnov statistic

Let 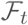 completely completely specify the realized history (σ-algebra) up to time *t* of a set of *n* observations indexed by *i* = 0, 1,…, *n* − 1 as specified in Section 2.1. Given this history, a researcher creates an event history model of both the event and censoring intensities for all observations *i*. Censoring will be interpreted as one event in a competing events framework. Let *φ_i_*(*t*) represent the comprehensive instantaneous intensity of competing events, Φ_*i*_(*t*) the associated cumulative intensity, and *R*_*i*_(*t*) = *e* ^−Φ*i*(*t*)^ the associated survival. We relax the assumption of independent censoring, requiring only that *R_i_*(*t*) be suitably modeled. The predicted martingale survival residual in terms of the comprehensive survival is

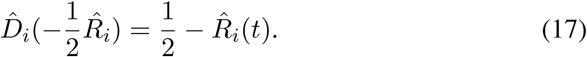

The probability density function and cumulative density function of the actual data residual for the comprehensive survival are

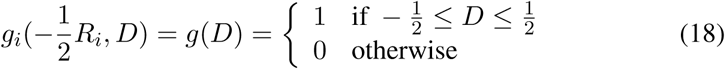
and

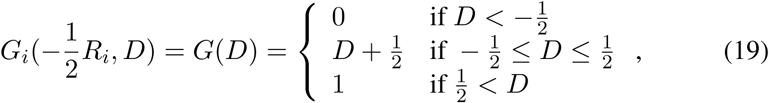

where *g*(*D*) and *G*(*D*) are the theoretical reference probability density function and cumulative density function, which are independent of the observation. This independence relies on the absolute continuity of *R*_*i*_(*t*). For a good model of the comprehensive transition intensities, the set of residuals 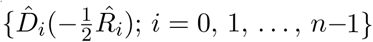 will be indistinguishable from a sample drawn from the reference uniform distribution, *g*(*D*), defined by Equation 18.

### 3.1 The Kolmogorov-Smirnov Statistic

To test whether the residuals are drawn from the uniform distribution, we use the Kolmogorov-Smirnov statistic. The Kolmogorov-Smirnov statistic is a sensible choice for the test because it is non-parametric, widely used, and has well-established statistical properties. The Kolmogorov-Smirnov test is distribution free if both the model and reference distributions are absolutely continuous. Future work could explore the use of other tests, such as the Anderson-Darling statistics. The Kolmogorov-Smirnov statistic is

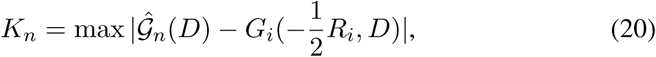
where

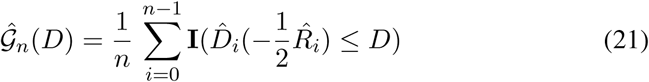
is the empirical cumulative distribution of the data residuals and **I**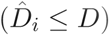 is the indicator function, equal to 1 if 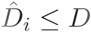 and 0 otherwise.

Well established procedures exist for calculating the statistical significance of the Kolmogorov-Smirnov statistic (i.e., the probability that a value larger than *K_n_* would be measured if the empirical distribution is in fact drawn from the reference distribution) under the aforementioned assumption of absolute continuity.

## 4 Illustrations

In this section, we utilize simulated data to demonstrate the utility of the theoretical material described in the preceding sections. For each example, we specify a model for the actual event and censoring intensities, generate observations using the actual model, specify a candidate maximum-likelihood model with which to fit the simulated data, solve for the parameters of the candiate model, calculate and plot the candidate residuals, and apply the goodness-of-fit test to assess whether the candidate model is acceptable. Table 1 summarizes the results for all examples.

**Table 1:** Summary of results for Illustrations.

| Example | Sample Size | Actual Model | Candidate Model | KS Stat. | KS Signif. | Reject Model |
| --- | --- | --- | --- | --- | --- | --- |
| 1. Constant intensities | 1000 | $\lambda_i = .05$<br>$c_i = .0125$ | $\hat{\lambda}_i = \hat{\lambda}$<br>$\hat{c}_i = \hat{c}$ | 0.0182 | 0.90 | No |
| 2. Linear intensities | 1000 | $\lambda_i = .05 + .005t$<br>$c_i = .0125$ | $\hat{\lambda}_i = \hat{\lambda}$<br>$\hat{c}_i = \hat{c}$ | 0.0921 | 8.6e-8 | Yes |
| 3. Power calculation | multiple | $\lambda_i = .05 + .005t$<br>$c_i = .0125$ | $\hat{\lambda}_i = \hat{\lambda}$<br>$\hat{c}_i = \hat{c}$ | multiple | multiple | N/A |
| 4. Additive Covariate | 2500 | Additive<br>[details in text] | Additive<br>Cox<br>Multip. | 0.014<br>0.0337<br>0.0485 | 0.71<br>7e-3<br>1.5e-5 | No<br>Yes<br>Yes |

## 4.1 Example 1: Constant transition intensities for the simulated data and model

This example uses simulated data for *n* = 1000 observations for which the actual transition intensities for both events and censoring are constant. The transition intensity for events is λ= 1/20 = 0.05 and that for censoring is *c* = 1/80 = 0.0125. The censoring probability is *ρ* = *c*/(λ + *c*) = 0.2 and the actual comprehensive transition intensity is *Φ* = λ + *c* = 0.0625.

The estimated maximum likelihood comprehensive transition intensity is 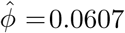, with an associated comprehensive survival of 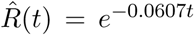. A histogram of the martingale survival residuals calculated using 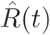 is plotted in Figure 4. The solid bar in the figure provides the theoretical probability density function as a reference. The Kolmogorov-Smirnov statistic for the data residuals is 0.0182 with a statistical significance of 0.8956. Since the empirical distribution of the data residuals cannot be distinguished from the theoretical reference distribution, we accept the model.

**Figure 3:**
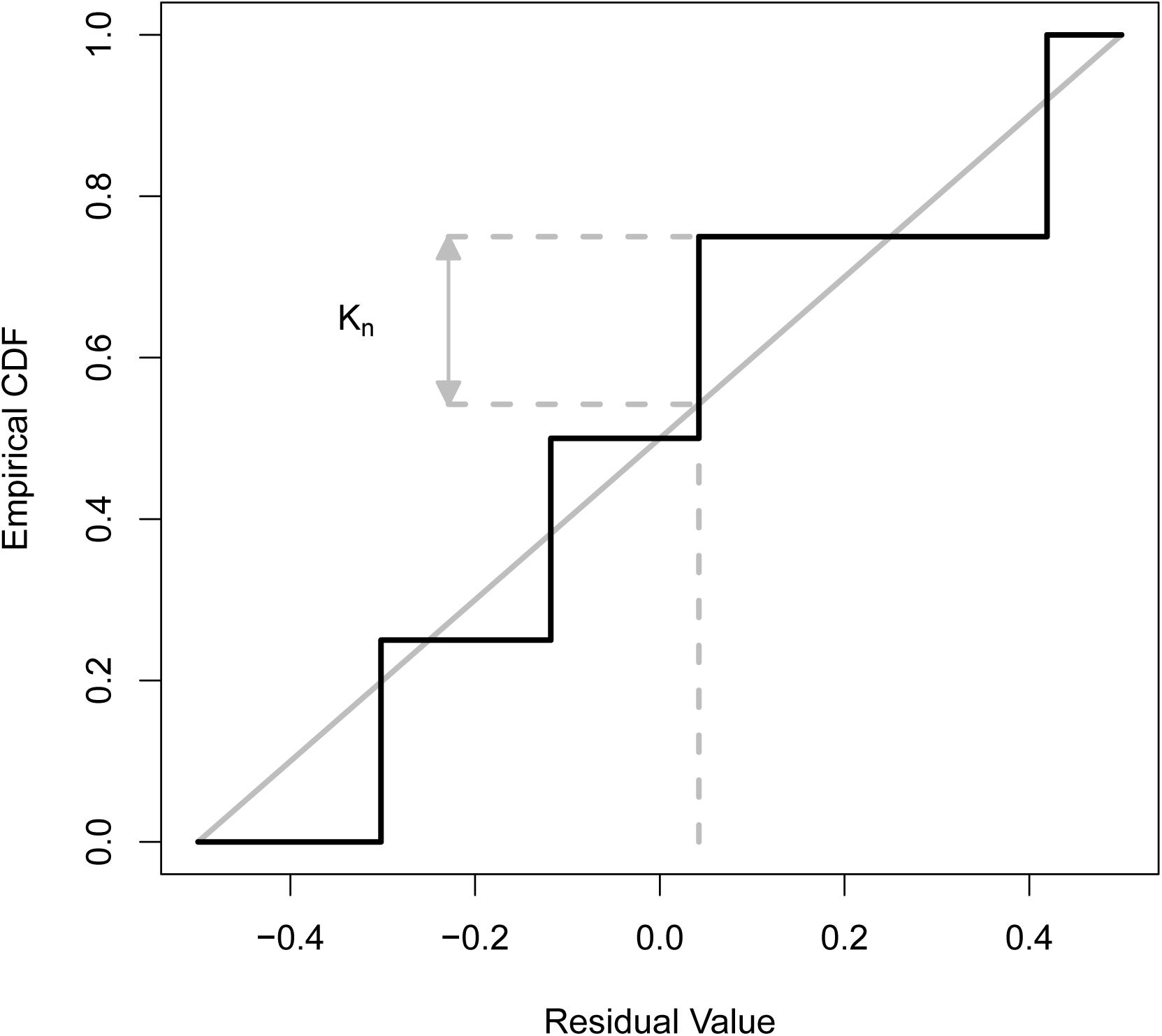
An empirical cumulative density function with four observations. The Kolmogorov-Smirnov statistics, *K_n_*, is the maximum distance between the empirical cumulative density function and the reference distribution (the non-dashed gray line).

**Figure 4:**
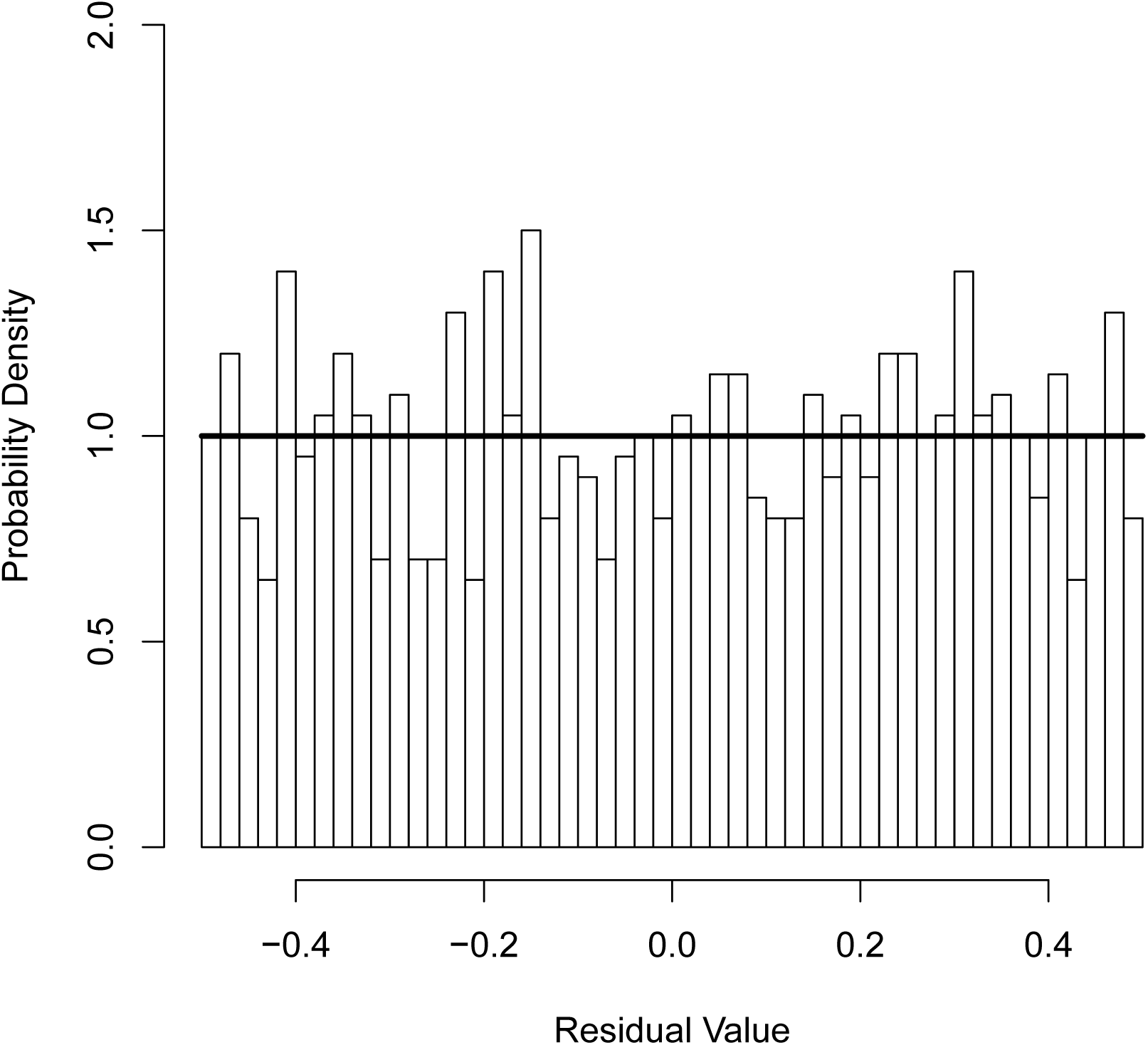
Example 1. Histogram of martingale survival residuals for a simula-tion with constant intensities of event occurrence and censoring (λ_*i*_ =.05 and *c_i_* =.0125). The percentage of censored observations given these parameters is 20%. 1000 simulated observations were created, a maximum likelihood estimate done assuming constant event and censoring rates (a correct model), and the data residuals calculated for each observation and displayed as a histogram. The theoretical residual distribution, which is a uniform distribution on the interval −0.5 to −0.5, is also shown. A Kolmogorov-Smirnov test indicates that the residuals do not differ from the uniform distribution (Table 1), so the proposed model is accepted.

### 4.2 Example 2: A mis-specified model

In this example, we deliberately mis-specify the model to establish that our goodness-of-fit test correctly rejects the mis-specified model. The transition intensity for events is λ_*i*_ =.05 +.005*t* (linear in time) and that for censoring is *c* = 1/80 = 0.0125 (constant; unchanged from the previous example). However, we incorrectly fit the data with the constant intensity model used in the previous example even though the intensity for event occurrence is linear in time.

A histogram of the martingale survival residuals calculated using 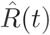 is plotted in Figure 5. The solid bar in the figure provides the theoretical probability density function as a reference. The mismatch between the data residuals and the reference distribution is visually clear in Figure 5, which is confirmed by the Kolmogorov-Smirnov test. The value of the statistic is 0.0921 with a statistical significance of 8.592 · 10^−8^. Since the empirical distribution of the data residuals can be distinguished from the theoretical reference distribution, we reject the model.

**Figure 5:**
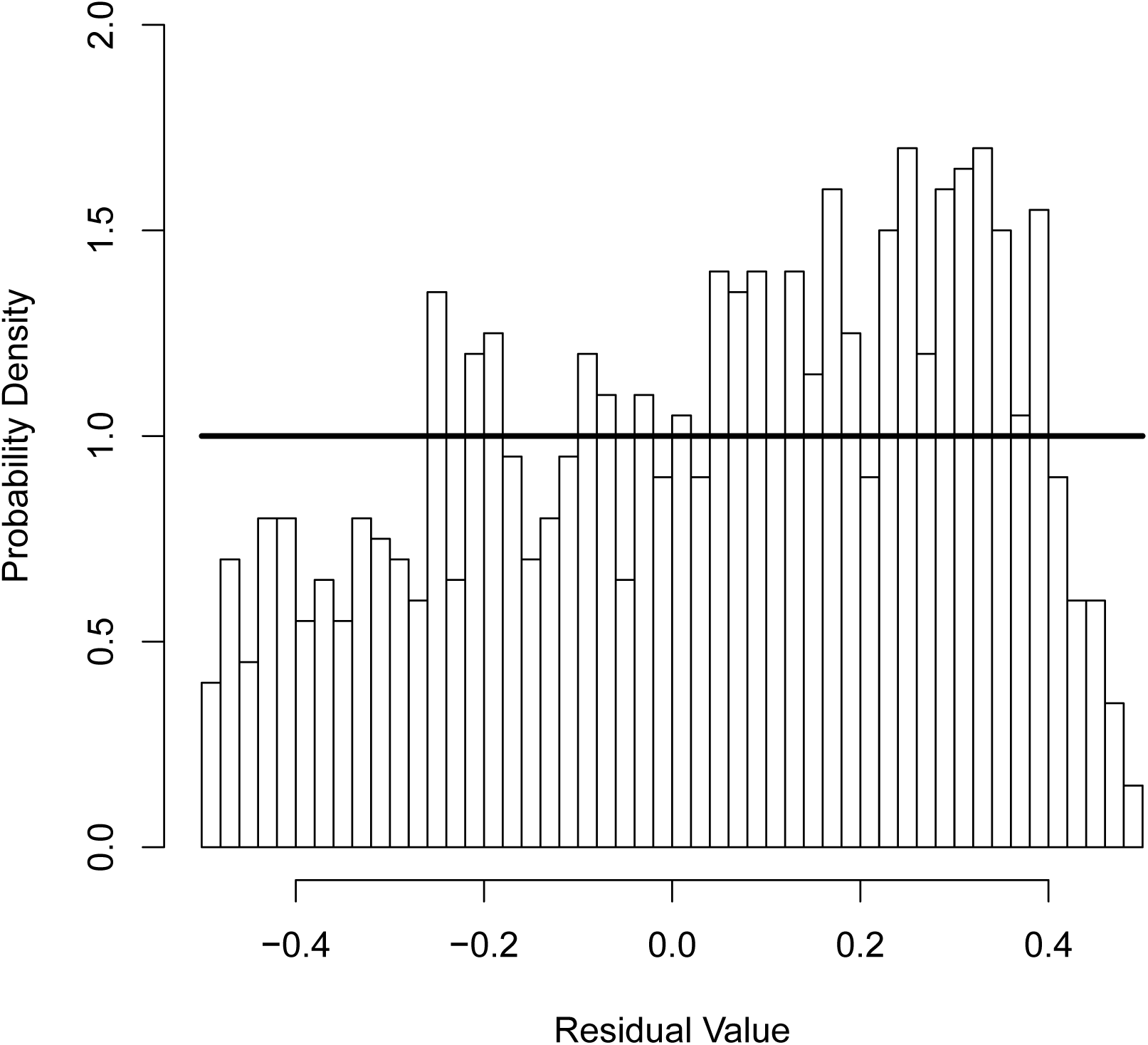
Example 2. Histogram of martingale survival residuals for a simulation with a linear, time dependent intensity for event occurrence (λ_*i*_ =.05 +.005*t*) and constant censoring (*c_i_* =.0125). 1000 simulated observations were created, a maximum likelihood estimate done assuming constant event and censoring rates (an incorrect model), and the data residuals calculated for each observation and displayed as a histogram. The theoretical residual distribution, which is a uniform distribution on the interval −0.5 to +0.5, is also shown. A Kolmogorov-Smirnov test indicates that the residuals do differ from the uniform distribution (Table 1), so the proposed model is rejected.

### 4.3 Example 3: A power calculation

In this example, we extend the previous example by conducting a power calculation using identical parameters, except that the number of observations is varied. The statistical power of a given test is the probability that the null hypothesis will be rejected when it is indeed false (Cohen 1992). Three factors determine the statistical power: (1) the significance criterion (*α*, usually 0.05); (2) the sample size (*n*); and (3) the effect size (*ES*). *α* is the criterion for Type 1 errors (mistakenly rejecting the null when it is true). The criterion for Type 2 errors is *β* (mistakenly accepting the null when it is false). The power is 1 − *β*.

We assume that *α* = 0:05 and that the effect size is *ES* = *b*_1_ = 0.005. The null hypothesis is that *b*_1_ is zero. Given these values, we calculate the power (1 − *β*) as a function of the sample size using 50,000 repetitions of the simulation. For each repetition, the test is successful if the p-value of the Kolmogorov-Smirnov test is less than *α* Figure 6 plots the power as a function of the sample size. The dashed lines show the x- and y-values of the curve for power levels of 0.8 (*β* = 0.2) and 0.9 (*β* = 0.1), two common cut-offs used in experiment design. The number of samples needed to achieve these power levels are 223 and 271, respectively. If the experimental sample size is lower than these values, the candidate model may be accepted even though it is incorrect. It is simply not possible to reject the model given the sample size.

**Figure 6:**
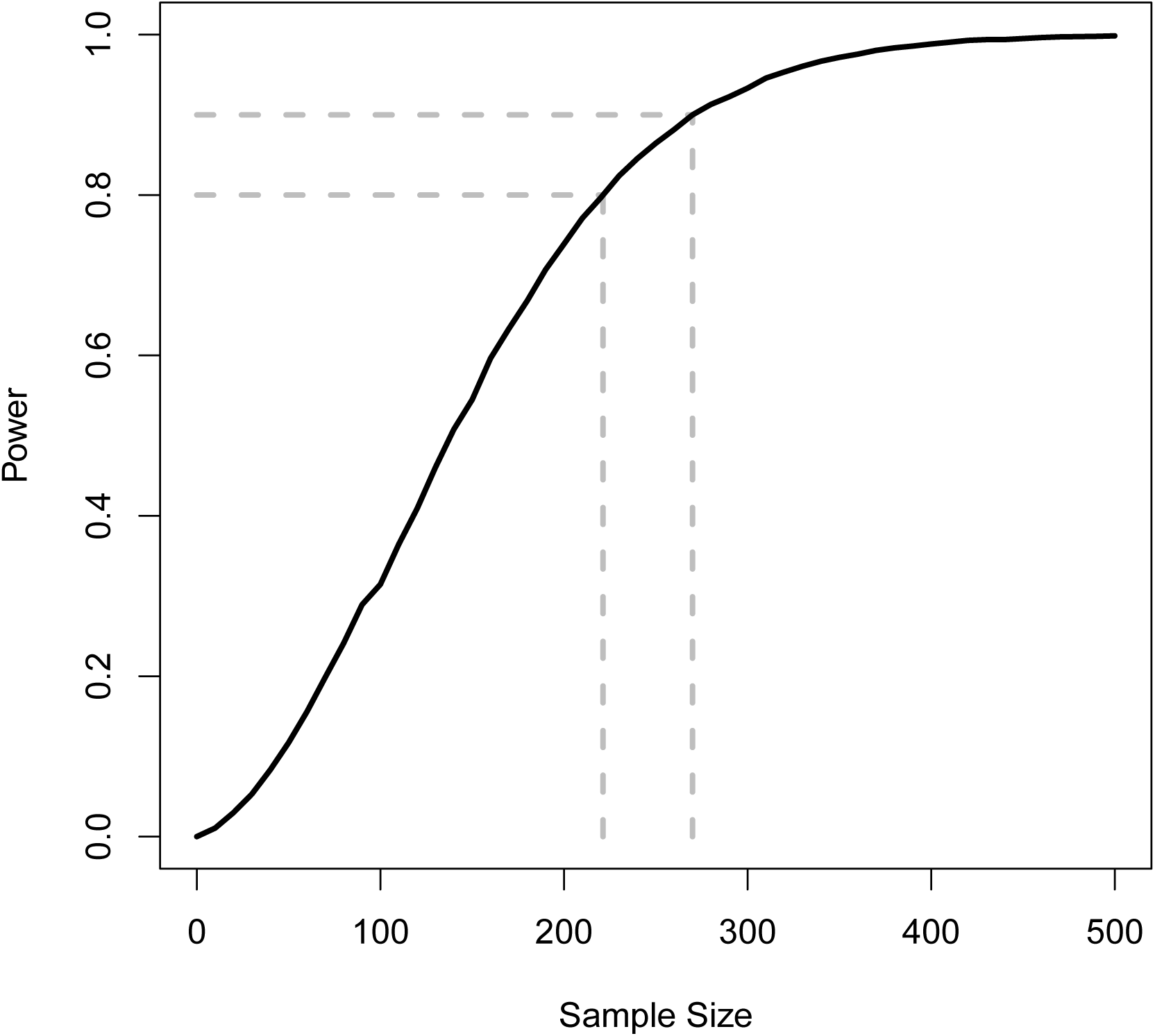
Example 3. Statistical power (y-axis) as a function of sample size (x-axis). The statistical power was calculated for sample sizes of *n* = 0, 25, 50, 75,…, 500. For each sample size *n*, 50, 000 repetitions of the simulation described in Example 2 were performed. The power is the number of simulations for each *n* for which the null hypothesis that *b*_1_ = 0 is correctly rejected at the *α* = 0.05 level. The effect size is the actual value of *b*_1_ used for the simulations, *b*_1_ = 0.005. The dashed gray lines show the samples sizes needed to achieve statistical powers of 0.8 (*β* = 0.2) and 0.9 (*α* = 0:1). The dashed gray lines intersect the x-axis at 222.2 and 270.9 so (rounding up) the sample sizes needed to achieve statistical powers of 0.8 and 0.9 are, respectively, 223 and 271.

### 4.4 Example 4: Another mis-specified model (proportional hazards)

A common assumption when modeling survival data is the proportional hazards assumption for the influence of covariates on the hazard (Cox 1972). In particular, the hazard is modeled as

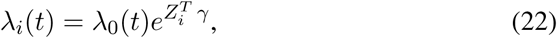

where λ_0_(*t*) is some baseline hazard, *Z_i_* is the vector of covariates for individual *i* (assumed here to be time-invariant), and γ is the vector of model coefficients that account for the magnitude of the influence of each covariate. However, in practice the proportional hazards is assumption is rarely tested. In this example, we simulate the situation in which the proportional hazards assumption is incorrect to demonstrate that our goodness-of-fit test can correctly reject the incorrect model. We model the true hazard as

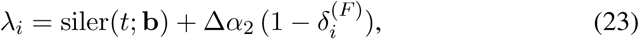
where

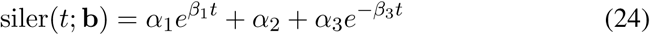
is the Siler hazard parametrized by the vector b = [*α*_1_;*β*_1_; *α*_2_; *α*_3_; *β*_3_], *δ_i_*^(*F*)^ is a covariate that is 1 if individual *i* is female and 0 if individual *i* is male, and Δ*α*_2_ is an additive boost to male mortality. Individuals are equally likely to be female and male. The values used to parametrize the Siler hazard are from a fit to demographic data from a cohort of women in Utah in the mid-1800s, b = [1.58 · 10^−1^; 1.53; 3.92 · 10 ^−3^; 4.83 · 10^−5^; 9.55 · 10 ^−2^]. The additional male mortality is Δ*α*_2_ = 5*α*_2_ = 0.020

We utilized these parameters to simulate age at death for 2500 individuals (1259 female and 1241 male). We then fit the sample of ages at death with three models: (1) the correct model (Additive, fully parametric), (2) Cox proportional hazards (Cox, semi-parametric), and (3) the following, incorrect multiplicative model (Multiplicative, fully parametric):

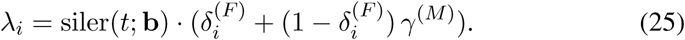

That is, the hazard for females is siler(*t*; b) and the hazard for males is proportional to this hazard, γ^(*M*)^ siler (*t*; b), where γ^(*M*)^ is the constant of proportionality. Figures 7 and 8, and 9 plot the martingale survival residuals for the three model fits. The values of the test statistic and statistical significance (in parentheses) are, respectively, 0.014 (0.71; additive), 0.0337 (7e-3; Cox), and 0.0485 (1.5e-5; multiplicative). Hence, the correct additive model is accepted, but both the Cox and multiplicative models are rejected per the test statistic.

**Figure 7:**
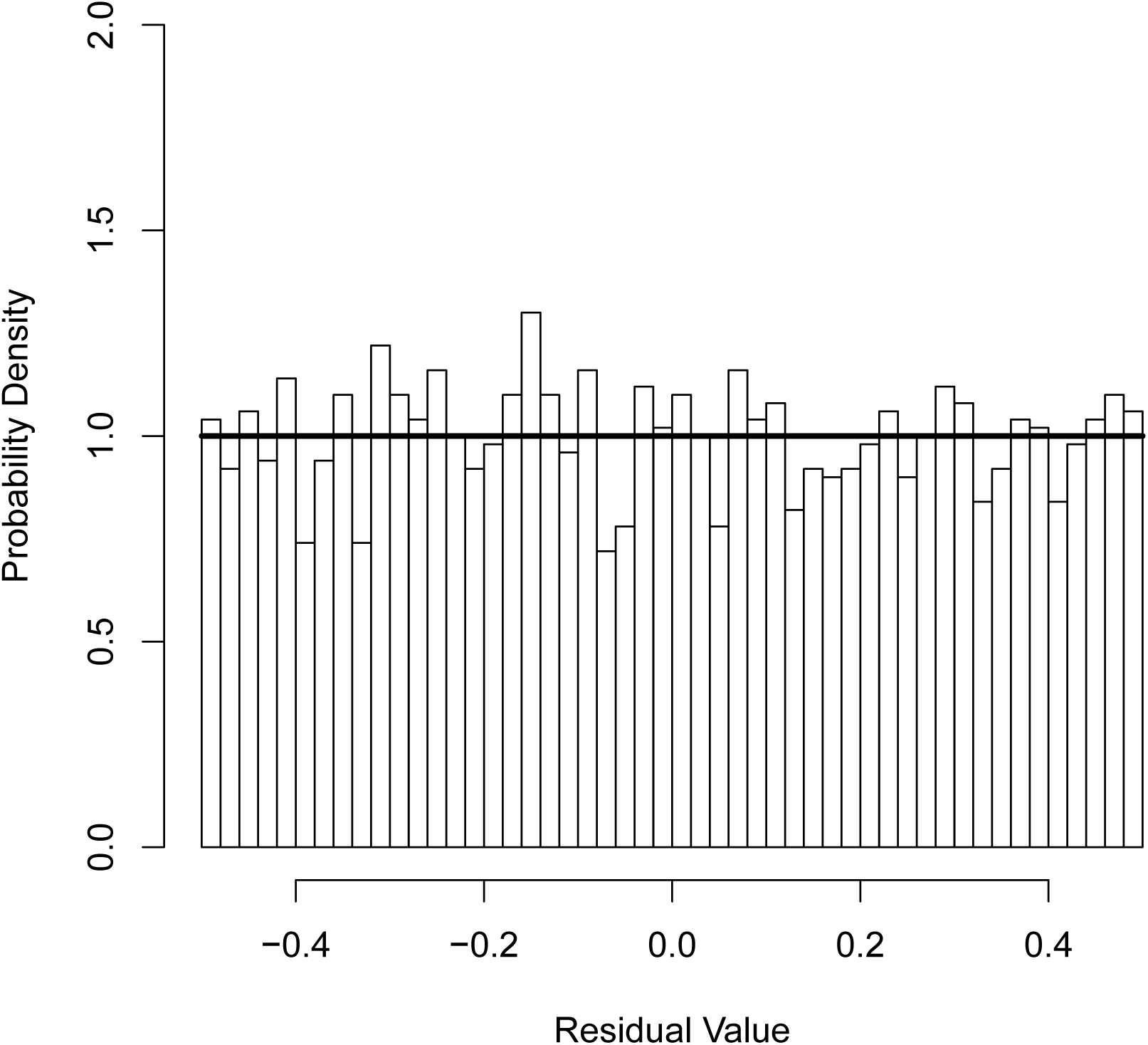
Example 4. Histogram of the martingale survivals residuals for the fully parametric additive fit (correct model) in the proportional hazards example. The Kolmogorov-Smirnov test accepts that the residuals are drawn from the uniform distribution (Table 1).

**Figure 8:**
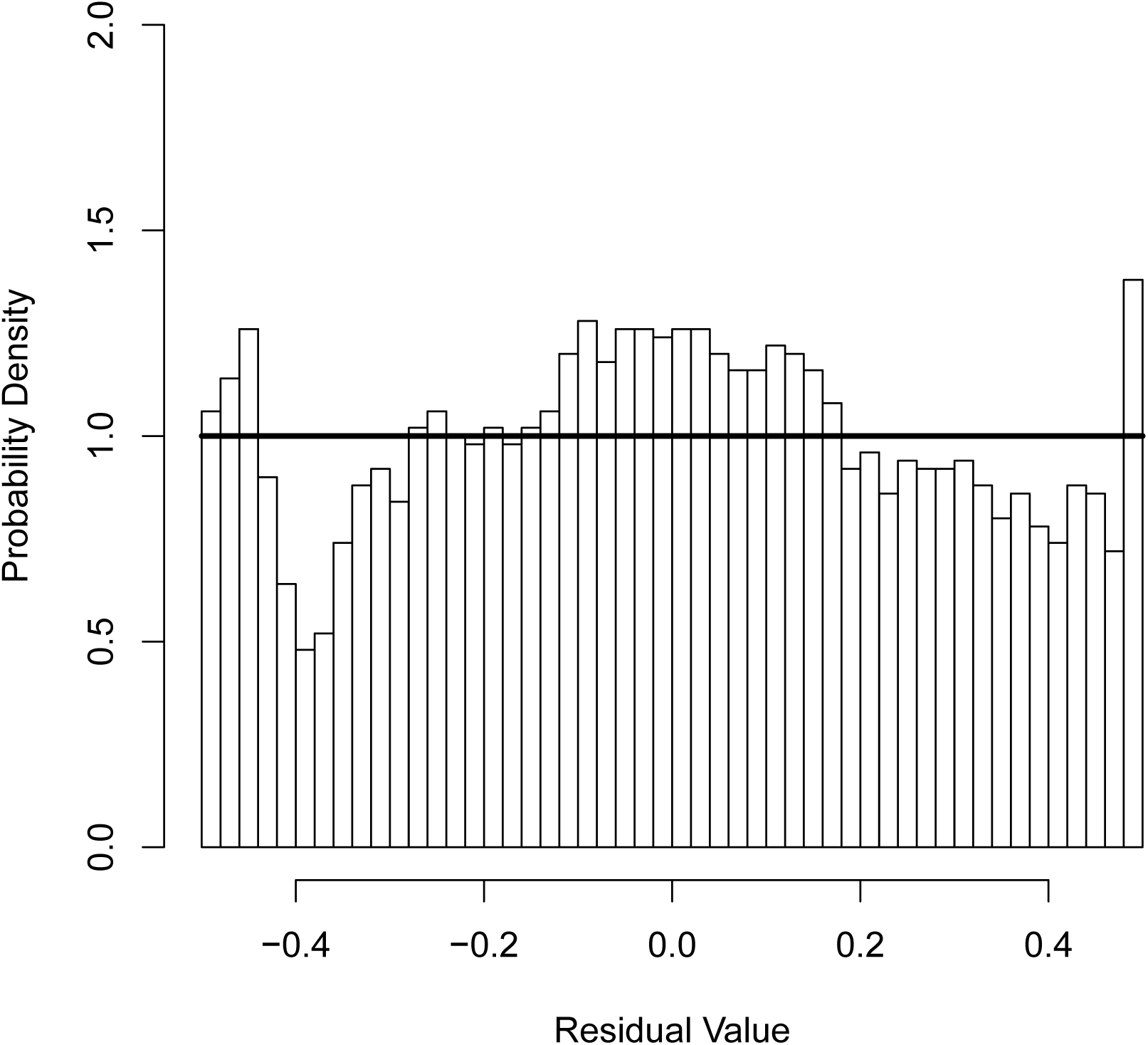
Example 4. Histogram of the martingale survivals residuals for the semi-parametric proportional hazards fit (incorrect model) in the proportional hazards example. The Kolmogorov-Smirnov test rejects that the residuals are drawn from the uniform distribution (Table 1).

**Figure 9:**
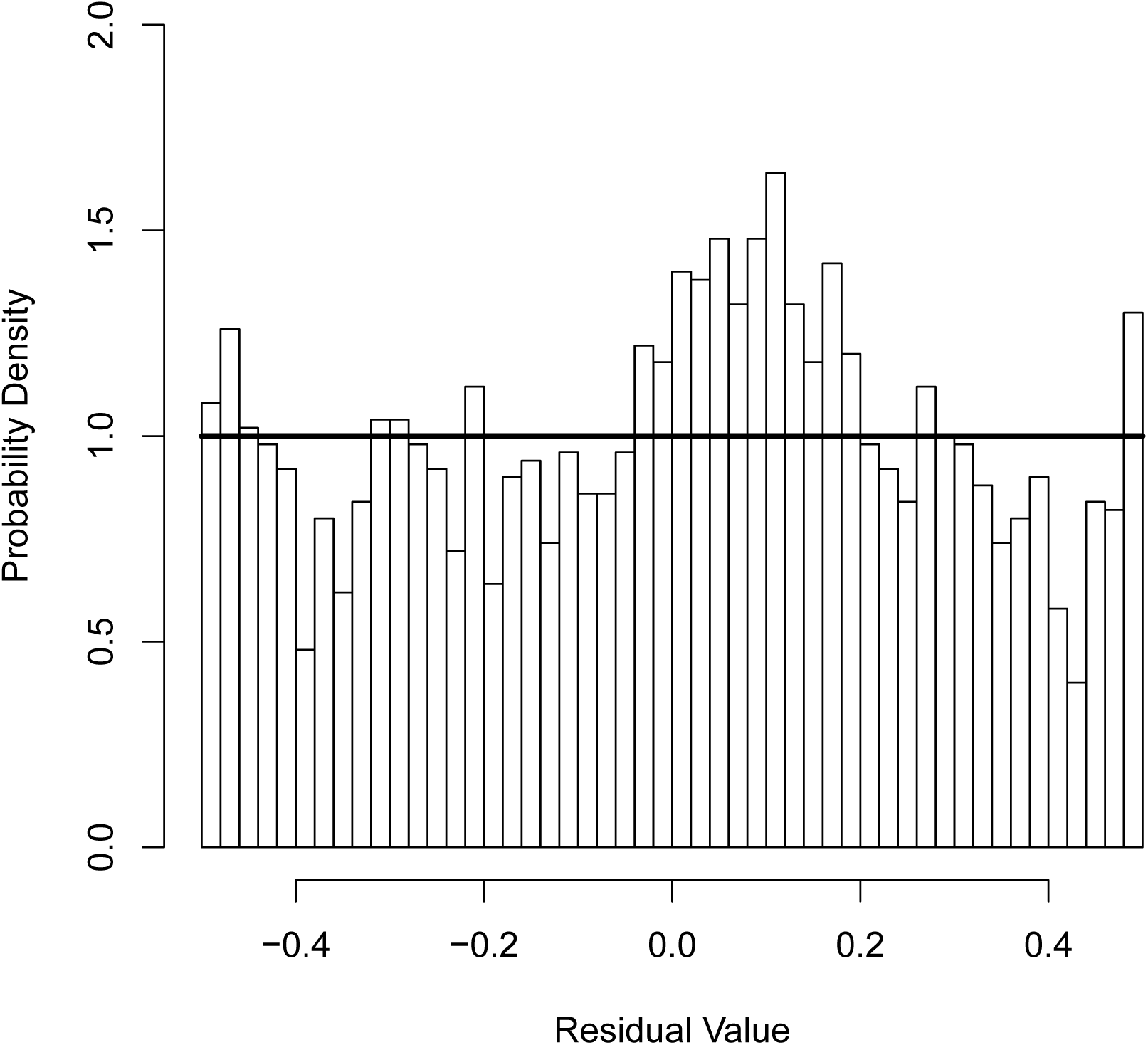
Example 4. Histogram of the martingale survivals residuals for the fully parametric multiplicative fit (incorrect model) in the proportional hazards example. The Kolmogorov-Smirnov test rejects that the residuals are drawn from the uniform distribution (Table 1).

## 5 Discussion

The four examples described in the preceding sections demonstrate conclusively that our proposed goodness-of-fit test successfully rejects incorrect models. This is the main contribution of this article, but it is not the only one. Another fundamental insight is that modeling the censoring process is essential to applying the goodness-of-fit test. We suggest that this is not a unique aspect of our approach, but rather is an aspect of survival data that must be accounted for by any goodness-of-fit test since the censoring process is an inherent component of the realized distribution of event times. Given this, additional consideration is needed of the assumption we made that the reference distribution’s cumulative density function is absolutely continuous, which is not true of an important class of studies.

It is common for studies to have a termination date when the study ends or when the data is “frozen” for analysis. For such studies, the modeled intensities for all observations are continuous up to the termination date, at which point there is a discontinuity in the censoring intensity since censoring instantly occurs at the termination date. For each modeled observation, the termination date corresponds to a unique termination value of the predicted data residual, so that the reference distribution is no longer a uniform distribution. Fortunately, such studies can be accommodated via an extension of the theory. This is a topic of ongoing investigation for us, but we will briefly summarize how this can be done.

We choose to call cumulative density functions that are absolutely continuous up to some pre-determined x-value where the transition is guaranteed to occur terminating distributions. It is possible to generalize the formula for the empirical cumulative density function in the same manner that the Kaplan-Meier estimate is used to account for the number of individuals at risk of an event. Observations that reach the terminating point (i.e., the value of the martingale survival residual at the termination date) are only informative up to that point, just as conventionally censored observations are only informative up to the censoring date. The Kolmogorov-Smirnov statistic is defined with respect to this generalization of the empirical cumulative distribution function.

One complication with this approach is that the theoretical distribution of the Kolmogorov-Smirnov measure derived from consideration of Brownian bridges is no longer valid. In fact, is not clear that an analytic result for the distribution of the measure can even be derived even for special cases of ensembles of terminating distributions. However, it is straightforward to numerically calculate the distribution in order to estimate the statistical significance of Kolmogorov-Smirnov test derived from the generalization of the empirical cumulative density function, which is the line of research we are investigating. For real data, this requires simulating the study repeatedly which, in turn, requires that a survival model be fit given the observed data. For observations that did not reach the terminating date in the observed data set but might reach it if the study is re-simulated, it is necessary to specify the values of any time-varying covariates beyond the actual event time.

A related issue is the impact of using the same underlying data to both fit the survival model and calculate the martingale survival residuals. This is directly analogous to the complication first discussed by Lilliefors (1967) caused by first estimating the mean and variance of a sample and then using the Kolmogorov-Smirnov test to determine if the sample is drawn from a normal distribution. The process of fitting data reduces the degrees of freedom in the data, which effectively regularizes it and, on the whole, reduces the expected distance between the empirical cumulative distribution function and the reference distribution. If this effect is ignored, the test is not stringent enough; samples drawn from an incorrect distribution are not rejected often enough.

In terms of our examples, correctly accounting for this fitting effect would reduce the number of samples needed to reject an incorrect model. Fortunately, ignoring the effect does not invalidate the conclusion to reject the models. The effect should be especially prominent for models with a large number of degrees of freedom, such as the semi-parametric Cox proportional hazard model in Example 4, for which the baseline hazard is estimated at large number of points. As with terminating distributions, this requires numerical simulations to account for the reduction in degrees of freedom caused by fitting the data and is a topic of active research for us. To characterize the fitting effect, we suggest that maximum likelihood estimate of the model parameters be used to re-simulate the experiment, a new model be fit to the re-simulated data, and the resulting Kolmogorov-Smirnov measure be calculated for repeated re-simulations. This provides an estimate of the distribution of the measure under the assumption that the actual model equals the fitted model. One can also assess the accuracy of this approach when an incorrect model is both used to fit the data and re-simulate the experiment, although of course one cannot know a *priori* with real data that the model is incorrect.

## 6 Conclusion

We have described a novel data residual for survival analysis called the martingale survival residual as well as a flexible, parameter-free goodness-of-fit test based on the martingale survival residual. We demonstrated the efficacy of the goodness-of-fit test using simulated data. The test provides an effective way to assess the accuracy of critical assumptions often made in survival analysis, such as the proportional hazards assumption, and is therefore a valuable contribution to researchers in a range of fields, including the natural, social, and medical sciences.

## A Derivation of probability density functions for residuals

In this appendix, we provide a formula for calculating the probability density functions of generalized martingale residuals (Equation 7), and derive probability density functions for the negative martingale residual (Equation 11) and martingale survival residual (Equation 12). Equations 11 and 12 demonstrate that censoring influences the value of generalized martingale residuals. Consequently, the censoring process must be explicitly accounted for in deriving probability density functions of generalized martingale residuals. Let *c_i_*(*t*) represent the instantaneous intensity of censoring events, 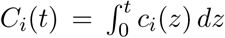 the associated cumulative intensity, and 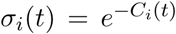 the associated survival.^1^ Let *φ_i_*(*t*) = λ*_i_*(*t*) + *c_i_*(*t*) represent the comprehensive instantaneous intensity, Φ*_i_*(*t*) = Λ*_i_*(*t*) + *C_i_*(*t*) the associated cumulative intensity, and *R_i_*(*t*) = *e*^Φ_*i*_(*t*)^ = *S_i_*(*t*) σ_*i*_(*t*) the associated survival. The multiplicative separability of *R_i_*(*t*) relies on the independence of events and censoring. Censoring can be interpreted as a competing event contributing to the comprehensive survival *R_i_*(*t*). The comprehensive cumulative density function as a function of time is

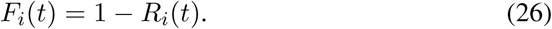

The corresponding probability density function is

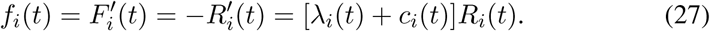

Re-integrating Equation 27, the cumulative density function can be split into a component due to event occurrence and a component due to censoring,

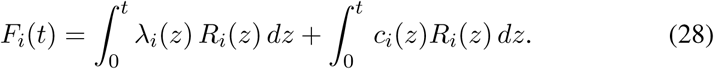

Equation 28 provides the cumulative density function as a function time, whereas what is desired is the cumulative density function as a function of the residual value. To calculate the latter cumulative density function, a change of variable from *t* to *D* can be applied to Equation 28. In general, separate mappings between *t* and *D* exist for non-censored and censored outcomes, which is why it is necessary to represent the cumulative density function in terms of the contribution of each. Let the functionals *u_i_*(*H_i_*, *D*) = *t* and *v_i_*(*H_i_*, *D*) = *t* represent the mappings for non-censored and censored outcomes, respectively, for the predictable process *H_i_*. ^1^

Utilizing Equation 28 and these mappings, the cumulative density function as a function of the observed residual *D_i_*(*H_i_*, *t_i_*) is

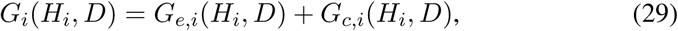
with

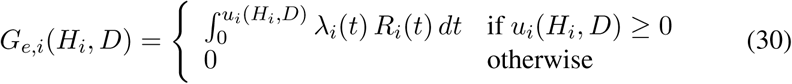
and

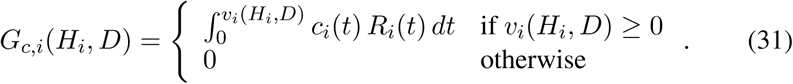

Recall that
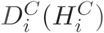 is the residual assuming no censoring (*C_i_*(*t*) = 0). The corresponding cumulative density function is

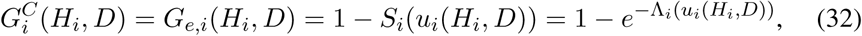
which relies on the fact that *R_i_*(*t*) = *S_i_*(*t*).

### A.1 Probability density functions of the negative martingale residual and martingale survival residual with no censoring

For the negative martingale residual (*H_i_* = −1), *u_i_*(−1, *D*) and *v_i_*(−1, *D*) can be found by solving Equation 11 for *t* with *δ_i_* equal to 1 for *u_i_*(−1, *D*) and 0 for *v_i_*(−1, *D*):

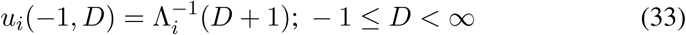
and

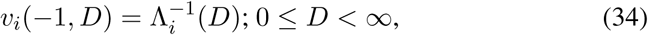

where the −1 in the exponent denotes an inverse.^2^ The cumulative density function and probability density function of the negative martingale residual with no censoring are

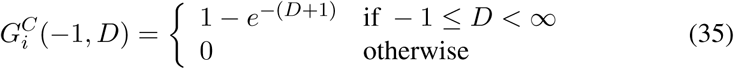
and

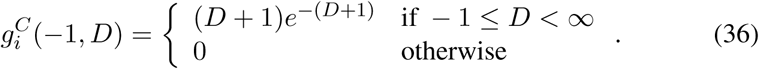

For the martingale survival residual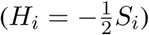, 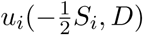 and 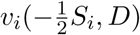 can be found by solving Equation 12 for *t* with *δ*_*i*_ equal to 1 for 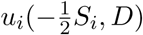 and 0 for 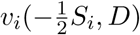:

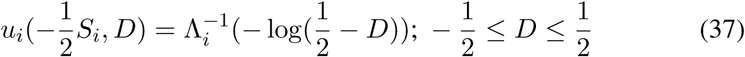
and

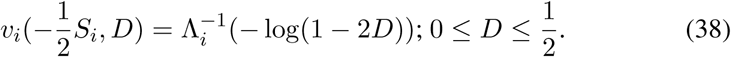

The cumulative density function and probability density function of the martingale survival residual with no censoring are and

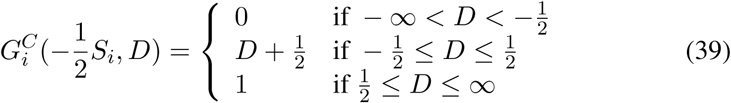
and

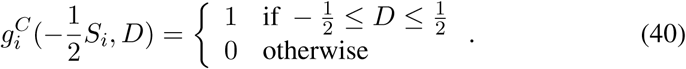

### A.2 Probability density functions of the negative martingale residual and martingale survival residual with constant transition intensities for event occurrence and censoring

If the transition intensities for event occurrence and censoring are both constant, Equations 30 and 31 simplify to

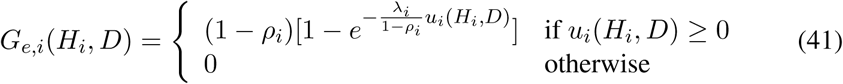
and

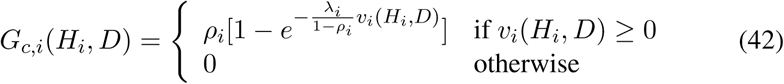

where *ρ*_*i*_ = *c*_*i*_/ (λ_*i*_ + *c*_*i*_) is the censoring ratio – that is, the probability that subject *i* is censored. The probability density functions for the negative martingale residual and martingale survival residual for constant transition intensities are, respectively,

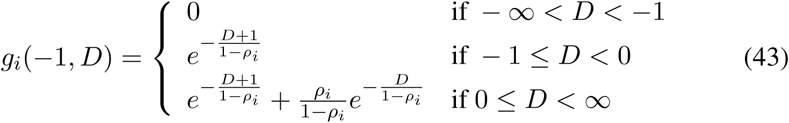
and

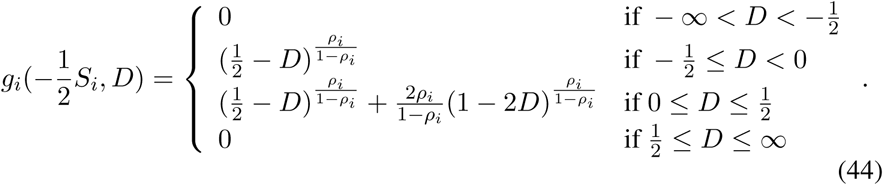

## Footnotes

1 There is an associated counting process for censoring, but we will not assign it a symbol since it is not explicitly used.

2 Since these derivations are for the probability density functions of the actual data residuals, no hats are needed.

